# Comparative *in vitro* antioxidant and antibacterial activities of leaf extract fractions of Crimson bottlebrush, *Callistemon citrinus* (Curtis.) Skeels

**DOI:** 10.1101/2021.03.26.436274

**Authors:** Poulomi Ghosh, Souren Goswami, Sujit Roy, Ria Das, Tista Chakraborty, Sanjib Ray

## Abstract

**Objectives:** The present study aimed to analyze a comparative *in vitro* free radical scavenging and antibacterial potentials of leaf aqueous and the successive extract fractions of *Callistemon citrinus*.

**Methods:** For *in vitro* antioxidant activity assessments, 1,1-diphenyl-2-picrylhydrazyl (DPPH) free radical scavenging activity, Fe^3+^ ion reducing antioxidant power (FRAP) assay, and total antioxidant capacity of the extracts were tested. Antibacterial potentials were tested through Agar well diffusion method using both gram-positive and gram-negative bacterial strains.

**Results:** Data indicate the polar successive aqueous fraction (AQF) possesses the highest free radicals scavenging capacity, with lowest EC_50_ (the required extract concentration to scavenge half of the free radicals) for DPPH and FRAP assays, and contains the highest total phenolics (308.2±5.9 µg tannic acid equivalent/mg dry extract, DE), flavonoids (516.7±3.5 µg quercetin equivalent/mg DE), total antioxidant capacity (441.48±12.8 µg ascorbic acid equivalent/mg DE). Phenolics and flavonoids contents were positively correlated with the *in vitro* antioxidant activities. The antibacterial study indicates the petroleum ether and chloroform are suitable solvents for extracting antibacterial phytochemicals from *C. citrinus* leaves that are effective against both gram-positive and negative bacterial strains.

**Conclusion:** The most polar fraction *i*.*e*. the successive aqueous extract fraction of *C. citrinus* leaves exhibited the highest antioxidant activities while the most non-polar petroleum ether extract fraction showed the highest antibacterial potentials thus these extract fractions might have therapeutic importance.

## 1. Introduction

In living cells, spontaneous production of a minimum level of free radicals is a normal physiological phenomenon, which is naturally maintained by *in vivo* antioxidant systems without causing any self-damage. This redox biology is very important to continue many physiological and biochemical actions, like cellular signaling in the host defense system^1^. The change in homeostasis between pro-oxidant and antioxidants leading to the pro-oxidant level high, moves towards oxidative stress. The reactive oxygen species (ROS), reactive nitrogen species (RNS), and their derivatives are produced in biological systems through the normal metabolic process and cause oxidative stresses while the endogenous counter mechanisms cannot balance those. Then we need to depend upon exogenous supplies through consuming antioxidant-rich foodstuffs or medicines to protect our body from their deleterious effects. Oxidative stress affects devastatingly on human health by inactivating the metabolic enzymes, damaging important cellular components, and oxidizing the proteins, nucleic acids, and lipids^2^,3. The standard antioxidant drugs like vitamin C, E, A, folic acid, gallic acid, etc. are mostly phenolics or flavonoids in nature. Antioxidants reduce the free radical burden of organisms by quenching them and thus they decrease the risk of oxidative stress-related pathologies^4^. Antioxidant quenches free radicals through several ways, like hydrogen-atom transfer (HAT) or single-electron transfer (SET) or both, and also repressing ROS/RNS generation by silencing some enzymes or chelating trace metals involved in reactive species production^1,2^. Several scientific reports indicated antioxidant properties of the different plant parts (leaves, fruits, barks or bulbs) and a strong positive correlation with their phenolics and flavonoids contents^5,6,7,8,9,10^.

It was reported that the microbial species have been taken up approximately 60% of Earth’s biomass, spread over nearly every habitat. Many of them are pathogenic and behind huge loss of human lives, livestock and crop productions. Overcrowded population is also a major reason to emerge and enhance in the infectious diseases over the last two decades. There are so many common broad and narrow-spectrum antibiotics such as amoxicillin, azithromycin, clindamycin, metronidazole, levofloxacin, tetracycline, kanamycin, etc. that are effective against both gram-negative and gram-positive bacteria, but these synthetic antibiotics show many side effects like increased oxidative stress, loss of appetite, diarrhea, indigestion or stomach pain^11^. Earlier reports are indicating that many plant extracts have rich antimicrobial as well as antioxidant properties^8,12^. Plant-derived natural compounds occupied a major portion of the pharmaceutical market just because of their fewer side effects, cost efficiency, and eco-friendly nature relative to the modern semi-synthetic or synthetic drugs.

*Callistemon citrinus* is an ornamental plant, mainly distributed in both the subtropical and tropical regions. It is commonly used in folk medicine to treat gastrointestinal pain, respiratory disorders, infectious diseases, etc.^13,14^. Scientific analysis revealed that this plant possesses antidiabetic, hypolipidemic, antioxidant^15^, cardioprotective^16^, hepatoprotective^17^, anti-*Helicobacter pylori*^18^, calcium channel inhibitor^19^, anti-elastase^20^, wound healing, antithrombic^21^, anticholinesterase and anti-inflammatory^15^ activities. Jamzad et al.^5^, reported a significant *in vivo* antioxidant activity of its flower extract, correlating with its flavonoids and phenolics contents. They also found that, its hydro-methanolic (1:4) extracts have a higher free radical scavenging potential than its essential oils. The essential oil content of this plant constitutes of some volatile aromatic and odoriferous compounds like 1,8-cineole, β-thujone, limonene, capillene, α-pinene, pulegone, β-pinene, etc. and are responsible for its bioactivities. Leaves and flowers of *C. citrinus* constitute a major compound, 1-(2,6-dihydroxy-4-methoxyphenyl)-3-methylbutan-1-one (phloroglucinol) which was reported for potent dose-dependent antinociceptive and anti-inflammatory properties correlating its strong *in vivo* antioxidant activity^22^. Different medicinal properties like thrombolytic, membrane-stabilizing, free radical scavenging and anti-inflammatory potentials of the different organic solvent-mediated leaf extracts of *C. citrinus* were studied and revealed to be very promising^23^.

Antibacterial potency of this plant was also evaluated. Its crude leaf ethanolic and 80% methanolic extract showed promising antimicrobial activity against gram-positive (*Bacillus anthracis, Bacillus cereus, Streptococcus pyrogenes*, and *Listeria monocytogenes*) and gram-negative (*Salmonella typhi, Klebsiella pneumoniae, Escherichia coli*, and *Pseudomonas aeruginosa*) bacterial strains^24^. Alkaloids, extracted from the leaves of *C. citrinus* were reported for bacterial growth inhibition with only 0.0025 and 0.835 mg/mL minimum bactericidal concentrations (MBC), respectively for *Staphylococcus aureus* and *Pseudomonas aeruginosa*^25^.

In the present state of knowledge, it is not clear about the antioxidant property of the crude leaf aqueous extract and the successive extract fractions of *C. citrinus*, and also their comparative *in vitro* antioxidant and antibacterial properties. Therefore, the present study was designed to analyze a comparative account of *in vitro* antioxidant and antibacterial activities of the leaf aqueous extract and the successive extract fractions of *C. citrinus* and to correlate these effects with their total phenolics and flavonoids contents.

## 2. Materials and methods

### 2.1. Chemicals

Glacial acetic acid and methanol were obtained from BDH Chemicals Ltd., UK. Ascorbic acid, DPPH, tannic acid, quercetin, and ferric chloride were purchased from Thermo Fisher Scientific Pvt. Ltd., Mumbai, India. Aluminum chloride, Muller Hinton agar, and Tetracycline were obtained from Himedia Laboratories Pvt. Ltd., Mumbai, India. Trichloroacetic acid (TCA) was purchased from Sigma Chemical Co, St.Louis, M.O. The USA. Sodium nitrite (NaNO_2_), sodium hydroxide (NaOH), sodium carbonate (Na_2_CO_3_), disodium hydrogen phosphate (Na_2_HPO_4_), dimethyl sulfoxide (DMSO) and Folin-Ciocalteu’s phenol reagent were obtained from Merck Specialities Pvt. Ltd., Mumbai, India. The used other chemicals were of analytical grade.

### 2.2. Plant collection, authentication, and storage

Fresh leaves of *C. citrinus* were collected from the Tarabag Residential Complex of The University of Burdwan. Prof. A. Mukherjee, Department of Botany, The University of Burdwan, has authenticated the plant taxonomically and a voucher specimen (No. BUR/TB/PG/01) is being preserved in the Department of Zoology. The collected leaves were washed thoroughly with tap water, shade dried, crushed into small pieces, made it into powder form with an electrical grinder (Philips Mixer blender HL 1605), and stored in an airtight glass container for future use.

### 2.3. Extracts preparation

20 g ground leaf of *C. citrinus* was extracted in 400 mL of distilled water at 80°C in a water bath and at every 4 h, the extract was collected and filtered through No. 1 Whatman filter paper. This process was repeated 3 times to get a large quantity of crude aqueous extract (CAQ) and it was then concentrated using a vacuum evaporator and stored at -20°C for future use.

Another 50 g of the leaf powder was successively extracted using a Soxhlet apparatus for 72 h with the different organic solvents (600 mL) of increasing polarity like petroleum ether, chloroform, ethyl acetate, methanol and distilled water at their boiling point. After every extraction, the remaining leaf powder was dried properly to remove the trace of organic solvents. All the extracts were then filtered through Whatman filter paper no. 1. and the successive extract fractions (SEF) [PEF; petroleum ether fraction, CHF; chloroform fraction, EAF; ethyl acetate fraction, MEF; methanol fraction, AQF; aqueous fraction] were dried completely using a rotary vacuum evaporator. Finally, these dried extract fractions were stored in airtight glass containers at 4°C for future use.

For *in vitro* antioxidant assays and quantitative phytochemical analysis the PEF, CHF, EAF, and MEF were dissolved in methanol, and AQF was dissolved in distilled water. For antibacterial assay PEF, CHF, and EAF were dissolved in 0.1% DMSO solution and MEF, AQF was dissolved in autoclaved distilled water.

### 2.4. *In vitro* antioxidant activity assessment

#### 2.4.1. DPPH free radical scavenging assay

The *in vitro* free radical scavenging activity of leaf aqueous extract and the SEF of *C. citrinus* leaves were determined through DPPH scavenging assay^26^. The standard ascorbic acid and different extract fractions (0.5 mL each) were taken in the respective test tubes and then 0.25 mL freshly prepared 1 mM DPPH methanolic solution and 1.5 mL of methanol was added to each. The test tubes were incubated for 35 min, in dark at room temperature (25±2°C) and then the optical density (OD) was measured at 517 nm using a UV-Vis spectrophotometer (UV-1800 Series, Shimadzu, Japan). A freshly prepared ascorbic acid solution of different concentrations (5-30 µg/mL) was used to prepare the standard curve. DPPH scavenging potential was calculated based on the following formula and the EC_50_ values (concentration at which 50% free radical scavenged), were also calculated.

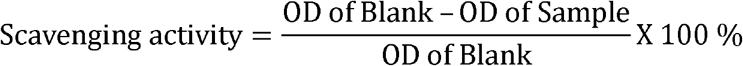

#### 2.4.2. FRAP assay

Another antioxidant assay, the ferric reducing antioxidant power assay is based on the transformation of ferric ion (Fe^3+^) to ferrous (Fe^2+^)^27^. 0.5 mL of extract, an equal volume of phosphate buffer (0.2 M), and 1% potassium ferricyanide [K_3_Fe(CN)_6_] were mixed well and incubated for 20 min at 50°C. Then 1 mL trichloroacetic acid (10%) was added to the mixture and centrifuged immediately for 10 min at 3000 rpm. Subsequently, in 1.5 mL of the supernatant, equal volume of distilled water and 0.1 mL of 0.1%, FeCl_3_ solution was mixed well. Then the OD was measured at 700 nm using a UV-Vis spectrophotometer. For standard curve preparation, freshly prepared ascorbic acid solutions of different concentrations (20-120 µg/mL) were used and the EC_50_ values for every fraction were calculated.

#### 2.4.3. Total antioxidant capacity (TAC) assay

The phosphomolybdate assay is widely used for TAC assessment of natural products^28^. The extract fractions (1 mL; 100 μg/mL) were mixed with the reagent mixture (4 mL; 0.60 M of sulfuric acid, 4 mM of ammonium molybdate, and 28 mM sodium phosphate) and incubated at 95°C for 90 min. The optical density was recorded at 695 nm using the UV-Vis spectrophotometer. The TAC of the extracts was calculated from the standard curve, prepared using ascorbic acid, and expressed as ascorbic acid equivalence (AAE).

### 2.5. *In vitro* antibacterial susceptibility test: agar well diffusion method

The *in vitro* antibacterial susceptibility test of the different extracts of *C. citrinus* was evaluated through the agar well diffusion method, based on the diameter of the zone of inhibition^29^. This *in vitro* microbial growth inhibitory activity of *C. citrinus* was determined against four human pathogenic bacteria, two of them were gram-positive such as *Bacillus megaterium* (MTCC 2412) and *Bacillus cereus* (MTCC 1272) and two were gram-negative such as *Staphylococcus aureus* (MTCC 9542) and *Enterobacter aerogenes* (MTCC 111). All bacterial strains were obtained as pure culture from the Parasitology and Microbiology Research Laboratory, Department of Zoology, The University of Burdwan, where they were maintained as stock strains in glycerol at -80°C.

For the antimicrobial assay, the pure cultures of the four above mentioned bacterial strains were inoculated separately on the sterilized nutrient broth media and incubated in BOD shaker incubator at 32±2°C temperature for 24 h. Sterilized plates containing Muller Hinton agar were prepared and 10 µL bacterial suspension was spread evenly over them, with a sterile cotton-glass hockey stick, from their standardized dilution of 10^8^ colony-forming unit (CFU)/mL of the microbes. Wells of 6 mm (diameter) were made with the help of Cork borer and from 1 mg/mL concentration of all the extract fractions 100 µL (≡ 100µg) were added in the separate wells. Tetracycline (30 µg/disc) was used as a positive control and DMSO (0.1 %) was used as a negative control. The plates were incubated in the BOD incubator for the next 24 h and then the zone of inhibitions was recorded from at least three of the replicas.

### 2.6. Phytochemical analysis

#### 2.6.1. Qualitative phytochemical analysis

The qualitative phytochemical analyses of SEF and the CAQ were done following the standard procedures as described by Harborne^30^ and Trease *et al*.^31^ with slight modifications.

#### 2.6.2. Quantitative analysis

##### 2.6.2.1. The total phenolics content (TPC)

The total phenolics content of the CAQ and SEF of *C. citrinus* leaves was estimated with Folin-Ciocalteu’s phenol reagent (FCP) and expressed as tannic acid equivalent (TAE) per mg of dried extract (DE)^32^.

At first, 100 µL (≡100 µg) of the test sample was taken in each test tube and then 900 µL distilled water was added to make the total volume of 1 mL. Then 500 µL of FCP (1 N) was added in each tube and mixed properly. After that, 2.5 mL of sodium carbonate (Na_2_CO_3_) solution (20%) was added and mixed thoroughly. Then they were kept in dark for 40 min at room temperature (25±2°C). After that, the OD was recorded at 725 nm using the UV-Vis spectrophotometer. The total phenolics content was estimated comparing the OD with the standard curve of tannic acid, prepared from varying range of concentrations (5-50 µg/mL). This assay was done in triplicate for each sample and the results were expressed as Mean±SEM.

##### 2.6.2.2. The total flavonoids content (TFC)

The total flavonoids content of all the extracts was estimated with aluminum chloride (AlCl_3_) method and expressed as quercetin equivalent (QE) per mg of dried extract (DE)^33^.

At first, 0.5 mL of the test sample was taken in each test tube and then 1 mL of distilled water was added. After that 1.5 mL of 5% sodium nitrite solution (NaNO_2_) and 0.15 mL of 10% AlCl_3_ were added. This mixture was allowed to settle down for 6 min and then 1 mL of 1M sodium hydroxide (NaOH) was mixed to all. After that water was added to the solution until the final volume reached 5 mL. Their OD was recorded at 510 nm using the above mentioned UV-Vis spectrophotometer, and from that, the flavonoids contents of the extracts were estimated comparing with the standard curve of quercetin. The range of quercetin concentrations for the standard curve preparation was from 10 µg to 500 µg/mL. This assay was also done in triplicate for each sample and results were expressed as Mean±SEM.

### 2.7. Statistical analysis

All data were expressed as Mean±SEM using Origin 8.0 Software and EC_50_ values were determined by statistical software. The significance of differences among the groups was calculated statistically by one-way analysis of variance (ANOVA) at the confidence level of 95% (*p*≤0.05) followed by the Post Hoc Tukey test using the Origin 8.0 Software. The correlation of coefficient (*r*) between the total phenolics and flavonoids content as well as total antioxidant caacity and also with the EC_50_ values of DPPH and FRAP assays were performed using Microsoft Office Excel 2007.

## 3. Result

### 3.1. *In vitro* antioxidant assessment

#### 3.1.1. DPPH assay

All the data indicate a positive correlation between the concentration of the extracts and their free radical scavenging activities. Here, the AQF showed the maximum free radical scavenging activity amongst all test extracts, which doesn’t vary significantly from the standard (*p*≤0.05) in one way ANOVA. Its EC_50_ value (43.88±3.07 µg/mL), the required concentration to scavenge half of the free radicals, is nearly twice than that of the standard ascorbic acid (22.42±0.74 µg/mL). All the extracts except PEF and CHF showed significant efficiency in free radical scavenging activity. The comparative EC_50_ values of DPPH scavenging activities of the remaining extracts revealed as 2.6, 3.4, and 3.7 times higher than the standard’s value respectively for EAF, MEF, and CAQ fractions (Table 1).

**Table 1:** Different *in vitro* antioxidant assays, total phenolics, and total flavonoids contents of the different extracts

| Extract<br>Fractions | DPPH assay<br>$EC_{50}$ values ($\mu\text{g/mL}$) | FRAP assay | TAC ($\mu\text{g}$<br>AAE/mg DE) | TPC ($\mu\text{g}$ TAE<br>/mg DE) | TFC ($\mu\text{g}$<br>QE/mg DE) |
| --- | --- | --- | --- | --- | --- |
| AA | $22.42 \pm 0.7$ | $28.54 \pm 2.3$ | - | - | - |
| PEF | $147.99 \pm 4.8$ | $653.19 \pm 5.8$ | $149.22 \pm 5.1$ | $37.37 \pm 6.3$ | $264.84 \pm 22.6$ |
| CHF | $512.02 \pm 14.3$ | $1188.21 \pm 11.8$ | $59.06 \pm 2.2$ | $36.85 \pm 2.1$ | $110.55 \pm 11.3$ |
| EAF | $58.92 \pm 0.9$ | $142.62 \pm 2.7$ | $259.24 \pm 6.9$ | $179.98 \pm 10.0$ | $232.95 \pm 13.9$ |
| MEF | $76.49 \pm 2.4$ | $182.50 \pm 2.4$ | $251.40 \pm 5.9$ | $207.40 \pm 6.8$ | $226.77 \pm 4.8$ |
| AQF | $43.88 \pm 3.0$ | $88.74 \pm 3.5$ | $441.48 \pm 12.8$ | $308.21 \pm 5.9$ | $516.70 \pm 3.5$ |
| CAQ | $83.28 \pm 3.4$ | $233.09 \pm 3.4$ | $241.33 \pm 1.1$ | $158.18 \pm 5.5$ | $193.13 \pm 3.3$ |

Data are expressed as Mean ± SEM. The level of confidence was considered at 95% ( $p \leq 0.05$ ) by ANOVA for the determination of differences among the groups followed by the Post hoc Tukey test. $EC_{50}$ - effective concentration to do the 50% effect; DPPH- 1,1-diphenyl-2-picrylhydrazyl; FRAP- ferric ion reducing antioxidant power; TAC- total antioxidant capacity; AAE- ascorbic acid equivalents; DE- dry extract; TPC- total phenolics content; TAE- tannic acid equivalents; TFC- total flavonoids content; QE- quercetin equivalents, PEF- petroleum ether fraction; CHF- chloroform fraction; EAF- ethyl acetate fraction; MEF- methanol fraction; AQF- aqueous fraction; CAQ- crude aqueous; SEM- standard error of the mean.

#### 3.1.2. FRAP assay

Like DPPH assay, the result of FRAP assay also indicates that AQF is the best depository of antioxidant phytochemicals. The half-maximum effective ferric ion reducing power concentrations (EC_50_) for standard ascorbic acid (28.54±2.3 µg/mL) was found to be one-third of the most effective fraction, AQF (88.74±3.5 µg/mL). The order of reduction capacity of ferric ion (Fe^3+^) to ferrous (Fe^2+^) of the remaining extracts were determined as EAF> MEF> CAQ> PEF> CHF, which reflects in their gradually increasing EC_50_ values respectively as 142.6±2.8, 182.5±2.5, 233.1±3.4, 653.2±5.8, 1188.2±11.8 µg/mL (Table 1).

#### 3.1.3. The total antioxidant capacity (TAC) assay

This data also indicates the differential TAC of the SEF and CAQ, and the decreasing order of TAC of them was established as AQF> EAF> MEF> CAQ> PEF> CHF, which is again similar to the previous order of EC_50_ values. The AQF showed a significantly higher (*P*≤0.05, analyzed by one way ANOVA, followed by Post Hoc Tukey test) antioxidant capacity (441.5±12.8 mg AAE/g dried extract) than all other extract fractions (Table 1).

### 3.2. *In vitro* antibacterial susceptibility test

For antibacterial activity assessment, the above-mentioned extracts were tested through the agar well diffusion method and found to have varying antimicrobial properties. Data designate the PEF as the most effective extract fraction against both the gram-positive and negative bacterial strains, followed by CHF and EAF. 29.0±1.7, 25.33±1.9, 23.0±1.2 and 22.0±1.5 mm zones of inhibition were formed by only 100 µg of PEF against *B. megaterium, B. cereus, S. aureus*, and *E. aerogenes* respectively (Table 2, Figure 1). The CHF (100 µg) showed 23.67±2.02, 24.33±1.8, 21±1.7 and 22.38±2.02 mm zones of inhibition against *B. megaterium, B. cereus, S. aureus* and *E. arogens* respectively. 15 to 17.33 mm zones of inhibition were made by the same dose of EAF against these bacteria. But other three extract fractions MEF, AQF, and CAQ didn’t have any antibacterial effect against all of the used bacterial strains. Also the negative control i.e. DMSO had not shown any zone of inhibition in any culture. The standard antibiotic tetracycline, which was used to compare the effectiveness of the extract fractions, had approximately 40 and 27 mm zone of inhibition against gram-positive and negative bacterial strains respectively.

**Table 2:** Antibacterial effects (zone of inhibition) of the test extracts, after 24 h of incubation

| Extract fractions | Diameter of zone of inhibition in mm (Mean±SEM) |  |  |  |
| --- | --- | --- | --- | --- |
|  | *B. megaterium* | *B. cereus* | *S. aureus* | *E. aerogenes* |
| CON-DMSO | 00.0±00.0 | 00.0±00.0 | 00.0±00.0 | 00.0±00.0 |
| TE | 40.67±0.88 | 39.0±2.08 | 26.33±2.02 | 27.60±1.76 |
| PEF | 29.0±1.73 | 25.33±1.85 | 23.0±1.15 | 22.0±1.52 |
| CHF | 23.67±2.02 | 24.33±1.76 | 21.0±1.73 | 22.33±2.02 |
| EAF | 17.0±1.15 | 15.0±1.52 | 17.33±2.33 | 17.33±2.33 |
| MEF | 00.0±00.0 | 00.0±00.0 | 00.0±00.0 | 00.0±00.0 |
| AQF | 00.0±00.0 | 00.0±00.0 | 00.0±00.0 | 00.0±00.0 |
| CAQ | 00.0±00.0 | 00.0±00.0 | 00.0±00.0 | 00.0±00.0 |

Data are expressed as Mean±SEM. The level of confidence was considered at 95 % ( $p \leq 0.05$ ) by ANOVA for the determination of differences among the groups followed by the Post hoc Tukey test. CON-DMSO= control dimethyl sulfoxide; TE= tetracycline; PEF= petroleum ether fraction; CHF= chloroform fraction; EAF= ethyl acetate fraction; MEF= methanol fraction; AQF= aqueous fraction; CAQ= crude aqueous; SEM= standard error of the mean.

**Figure 1:**
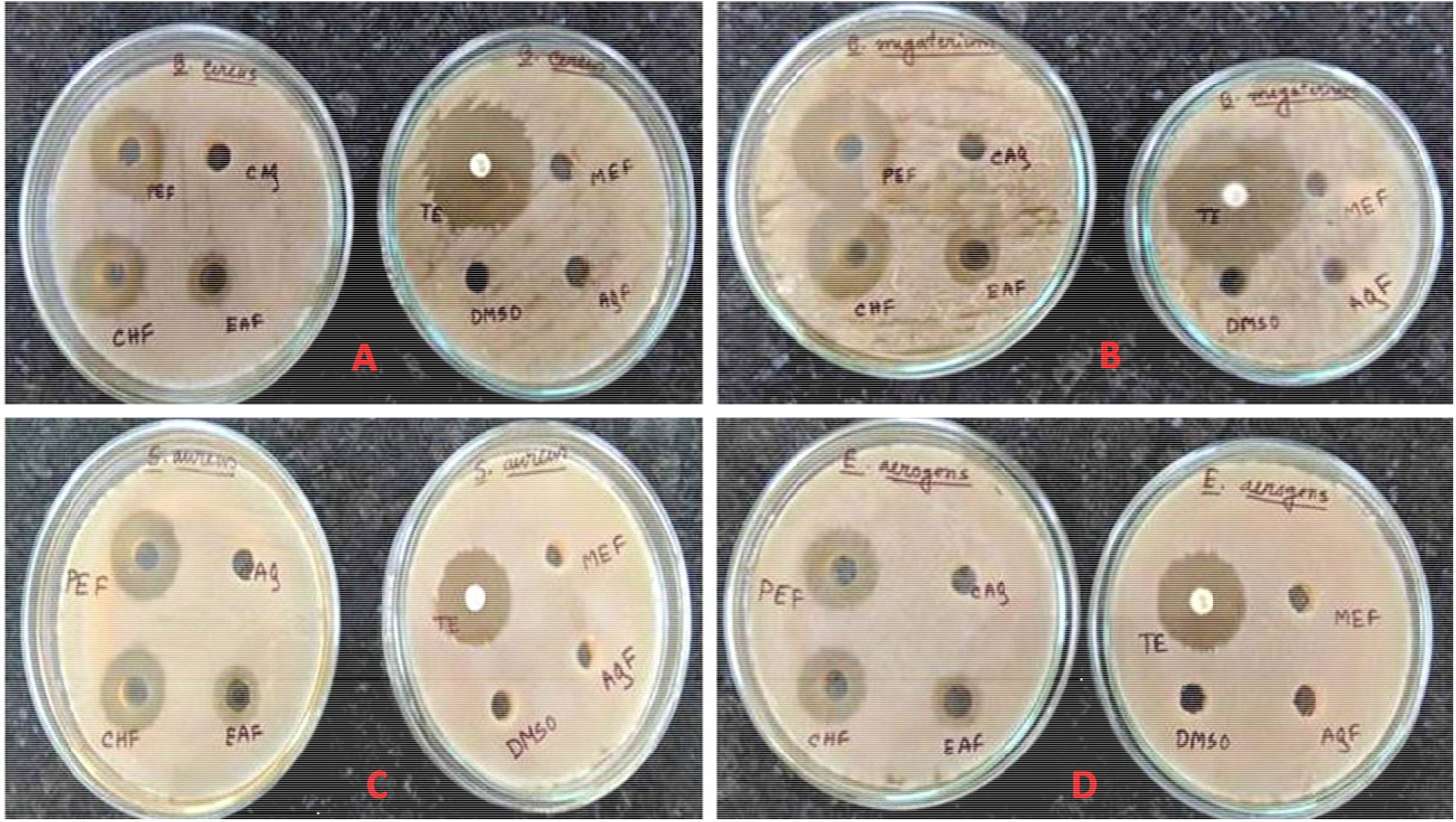
Photographs showing the zone of inhibitions with the successive extract fractions of *C. citrinus* leaves at 24 h of incubation against gram-positive and gram-negative bacteria. A; *Bacillus cereus*, B; *Bacillus megaterium*, C; *Stephylococcus aureus*, D; *Enterobacter aerogenes*. PEF= petroleum ether fraction; CHF= chloroform fraction; EAF= ethyl acetate fraction; MEF= methanol fraction; AQF= aqueous fraction; CAQ= crude aqueous; DMSO= dimethyl sulfoxide; TE= tetracycline.

### 3.3. Phytochemical analysis

#### 3.3.1. Qualitative phytochemical analysis

The preliminary phytochemical analyses show the presence of different types of secondary metabolites such as alkaloids, flavonoids, terpenoids, tannins, steroids, carbohydrates, and saponins in varying amounts, but glycosides, phlobatannins, and anthraquinones were absent in any of the extract (Table 3). Steroids were detected in PEF, CHF, and EAF. Terpenoids were detected in all fractions and found comparatively more abundant in MEF, AQF, and CAQ than the others, moderate in PEF and EAF, and detected least in CHF. Alkaloids were detected in PEF, CHF, and MEF. Flavonoids were detected in PEF, EAF, MEF, AQF, and CAQ.Tannins were detected in EAF, MEF, AQF, and CAQ. Carbohydrate and reducing sugars were detected only in the CAQ, MEF and AQF while saponins were detected in CAQ, and AQF.

**Table 3:**
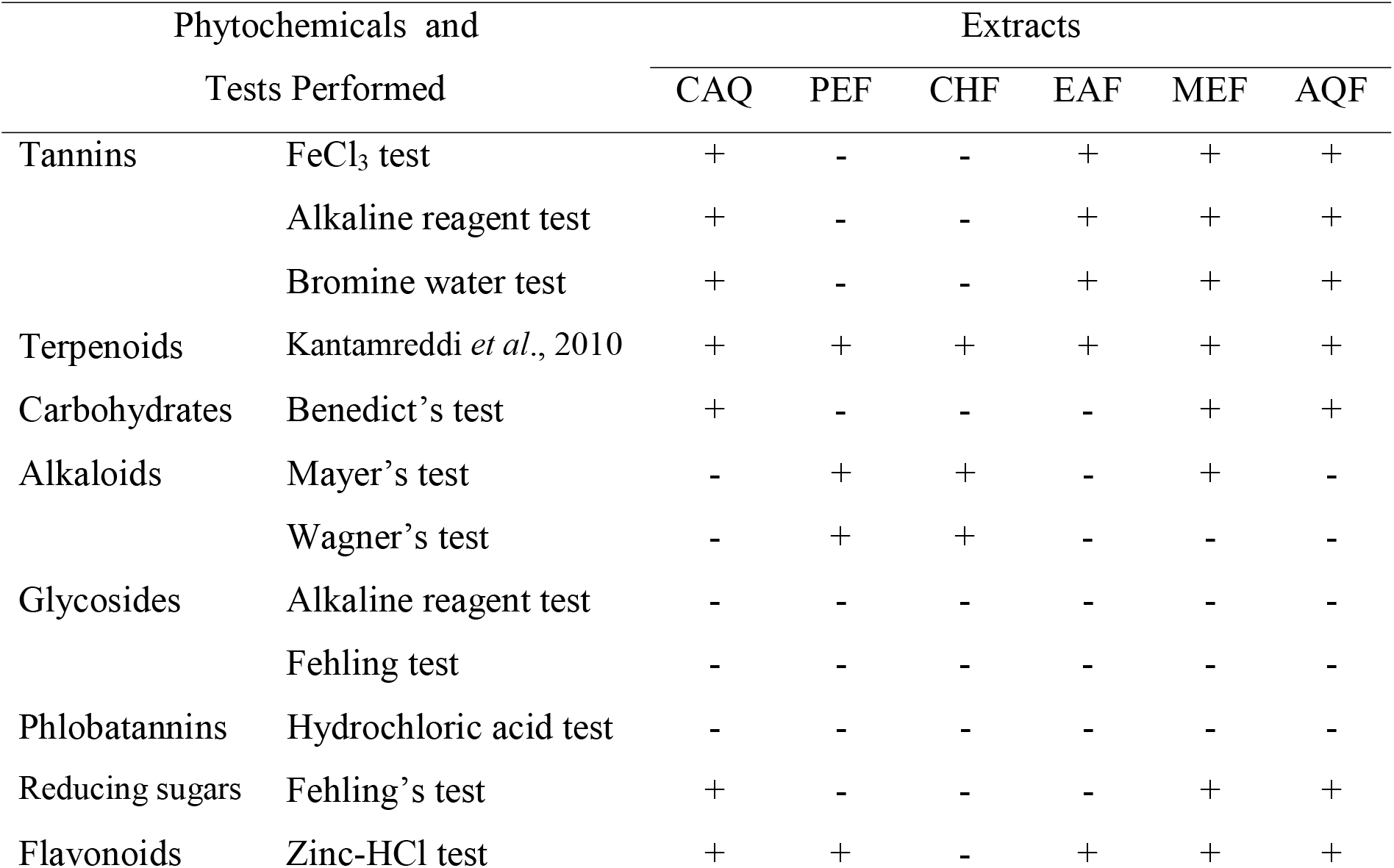
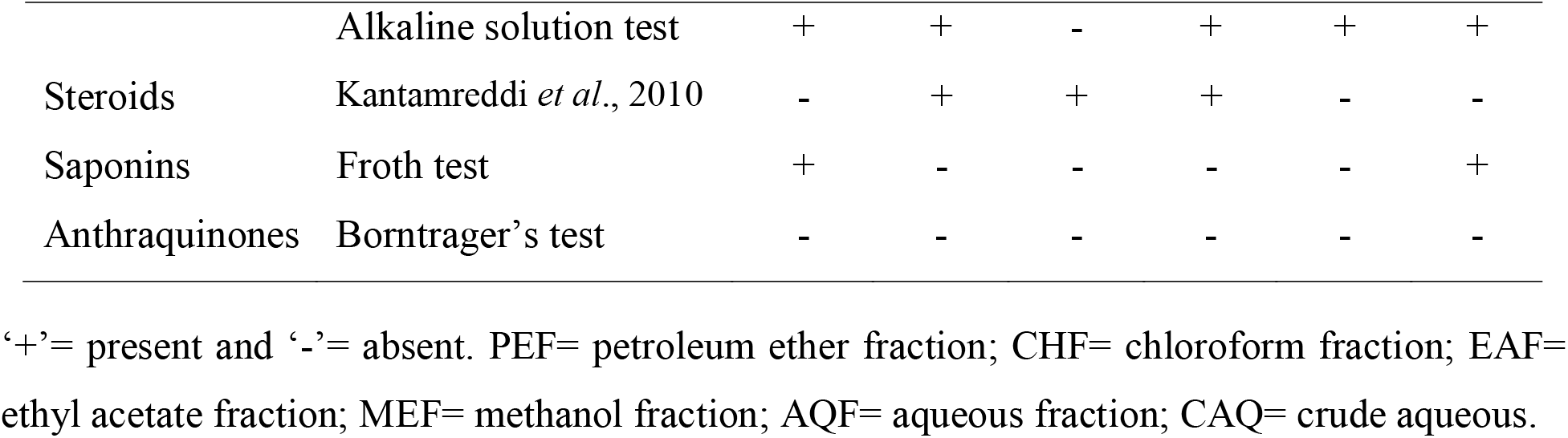
The preliminary phytochemical analysis of the test extracts

| Phytochemicals and Tests Performed |  | Extracts |  |  |  |  |  |
| --- | --- | --- | --- | --- | --- | --- | --- |
|  |  | CAQ | PEF | CHF | EAF | MEF | AQF |
| Tannins | FeCl<sub>3</sub> test | + | - | - | + | + | + |
|  | Alkaline reagent test | + | - | - | + | + | + |
|  | Bromine water test | + | - | - | + | + | + |
| Terpenoids | Kantamreddi *et al.*, 2010 | + | + | + | + | + | + |
| Carbohydrates | Benedict's test | + | - | - | - | + | + |
| Alkaloids | Mayer's test | - | + | + | - | + | - |
|  | Wagner's test | - | + | + | - | - | - |
| Glycosides | Alkaline reagent test | - | - | - | - | - | - |
|  | Fehling test | - | - | - | - | - | - |
| Phlobatannins | Hydrochloric acid test | - | - | - | - | - | - |
| Reducing sugars | Fehling's test | + | - | - | - | + | + |
| Flavonoids | Zinc-HCl test | + | + | - | + | + | + |
|  | Alkaline solution test | + | + | - | + | + | + |
| Steroids | Kantamreddi *et al.*, 2010 | - | + | + | + | - | - |
| Saponins | Froth test | + | - | - | - | - | + |
| Anthraquinones | Borntrager's test | - | - | - | - | - | - |

‘+’= present and ‘-’= absent. PEF= petroleum ether fraction; CHF= chloroform fraction; EAF= ethyl acetate fraction; MEF= methanol fraction; AQF= aqueous fraction; CAQ= crude aqueous.

#### 3.3.2. Quantitative analysis

##### Total phenolics and flavonoids contents

The total phenolics and flavonoids contents of the extract fractions of *C. citrinus* were estimated and data indicate that AQF contains the highest quantity of phenolics (308.2±5.9 µg TAE/mg DE) and flavonoids (516.7±3.5 µg QE/mg DE) (Table 1). The differential total phenolics contents of the extracts are in a decreasing order of AQF> MEF> EAF> CAQ> PEF> CHF and in the case of total flavonoids, that slightly differs with placing PEF fraction at the second-highest position rather from the second-lowest. The CHF contains the least quantity of both the phytochemicals.

## 4. Discussion

*Callistemon citrinus* has gained much attention as a folk medicinal reputation for the treatment of various ailments like gastrointestinal pain, respiratory disorder, infectious diseases, etc. and its various beneficial activities including antidiabetic, cardioprotective, and hepatoprotective potentials were explored scientifically^14,15,16,17,24,25^. In this study, we have explored a comparative *in vitro* antioxidant and antibacterial activities of the crude aqueous and the successive leaf extract fractions. Thus different *in vitro* free radical scavenging activities like total antioxidant, DPPH and FRAP assays, phytochemicals analysis, along with antibacterial tests were performed.

Out of the six extracts tested here, the last and first fractions of the Soxhlet extraction, i.e. the AQF and PEF have displayed the highest antioxidant and antibacterial activities respectively. The data of DPPH and FRAP assays indicate that the last Soxhlet fraction, AQF, is the best depository of antioxidant phytochemicals amongst all the extract fractions. Though the EC_50_ values of ascorbic acid for both DPPH and FRAP assays were lower than the most effective fraction, AQF, but despite being a mixture of compounds, its required concentration was only thrice to the standard. It is reported that most of the antioxidant compounds responsible for plants bioactivity are mainly phenolics and/or flavonoids which are extracted in polar solvents as the crude extract contains a rich combination of phytochemicals, while a specific fraction of any successive extraction procedure have fewer types of molecules due to their solvent affinity and polarity^6,8,9,10,28^. The differential total antioxidant activity (TAC) of the different organic solvent-mediated extracts of various medicinal plants were reported due to their variations in phenolics content^8,9,28^. Here also it is clear that the last fraction of the successive extraction procedure, which acts as the most potent antioxidant fraction, is rich in bioactive phytochemicals and extracted with the most polar solvent. An earlier study indicated that the crude leaf methanolic extract of *C. citrinus* and its extract fractions have differential free radical scavenging potentials and that also supports our findings that water is more suitable than petroleum ether for extracting bioactive antioxidant molecules from *C. citrinus* leaf^23^. Recently, through a comparative study, our laboratory reported that the successive leaf aqueous extract fraction of *Crinum asiaticum* having the highest antioxidant and antibacterial potentials along with the highest phenolics contents^8^.

From over 8000 known flavonoids molecules, polyphenols with C_6_-C_3_-C_6_ skeleton, have shown many *in vitro* multiple biological activities like anticancer, anti-oxidation, anti-inflammation, cardiovascular, and antibacterial^34^. Flavonoids and tannins which are phenolics were more or less detected in EAF, MEF, and AQF fractions, which have prominent free radical scavenging and antibacterial actions. There is also other evidence that crude methanolic and ethanolic extracts have more potency to scavenge free radicals than other crude extract or successive extract fractions^5,21^. Another study reported that ketones, terpenoids, aldehyde, hydrocarbons, and other groups also have minor contribution to quench free radicals. Several recent scientific reports indicate that antioxidant activity of various plant extracts are remarkably high and may health-beneficial for us^6,8,10,35,36^. We need to consume a larger amount of antioxidant-rich foods, such as different types of vegetables (pepper leaf, oregano, beans, beetroot, color cabbages, sweet potato, broccoli, carrots, etc.) and various fruits (berries, peach, banana, grapes, apple, pear, etc.), and beverages (tea, coffee, and various fruit juices, etc.) to reduce the probability of deadly diseases related to oxidative stress^35,37^.

The AQF of *C. citrinus* leaves shows a significantly higher amount of both TFC and TPC than other fractions, which correlates its maximized antioxidant activity and may be reasoned behind it. The TAC of AQF is also the highest of all. Plants produce various types of secondary metabolites such as flavonoids, alkaloids, steroids, terpenoids, etc., which protect plants from microorganisms, herbivores, insects or even other competitive plants that’s why a potential antibacterial activity is expected from such extracts. Quantitative phytochemical analysis revealed that PEF contains the second-highest total flavonoids amongst all the extract fractions. Flavonoids under the group phenolics, derived from tyrosine, phenylalanine, and malonate are well-known antioxidants. The chemical structure of phenolic compounds is ideal for free radical quenching due to the presence of hydroxyl groups with aromatic rings. Hydrogen atom or an electron of the hydroxyl group is easy to donate to a free radical and thus creates phenoxy radical intermediates, which are relatively stable than free radicals due to the resonance formation within the benzene ring^38^. There is sufficient literature regarding flavonoids’ antibacterial mechanism of action^39,40^. It is reported that flavonoids coat the cells’ surfaces and consequently disturb the interactions between the surface of the substratum and bacterial cells^41^. The antibacterial activity of flavonoids can be exerted by killing bacteria directly, synergistically activating antibiotics, and attenuating the bacterial pathogenicity through the combination of inhibitory actions on the nucleic acid synthesis, energy metabolism, cytoplasmic membrane function, attachment and biofilm formation^40^, restrained the synthesis of peptidoglycan and ribosome in the cells of amoxicillin-resistant *E. coli*^42^, inhibiting the efflux pump of MRSA^43^, inhibiting different kinds of lactamases, key enzymes that disable antibiotics, produced by bacteria^44,45^.

Many scientists explored the antibacterial activity of crude methanolic and ethanolic leaves extract, but here we found that the antibacterial compound(s) were extracted mainly in the non-polar solvents (petroleum ether, chloroform, and ethyl acetate) when successive extraction was done using a Soxhlet apparatus. Phytochemical tests showed that PEF, CHF, and EAF contain steroids, alkaloids, terpenoids, and flavonoids. The Steroids were comparatively more abundant in CHF than PEF but both of them were positive for terpenoids and alkaloids too. It was reported that *E. coli* is resistant to methanolic and ethanolic extracts of *C. citrinus* and its probable reason is cell membrane permeability or genetic factors^24^, this report supports the present study, all bacterial strains were resistant to MEF, AQF, and CAQ.

Many studies have revealed that the gram-positive bacteria are more susceptible than gram-negative bacteria due to the absence of gram-negatives’ unique outer structure i.e. outer membrane, periplasmic space, high lipopolysaccharide content, high lipid, and lipoprotein content^24^. Both the PEF and CHF show greater efficacy against gram-positive strains than the gram-negative, indicating the effect may be related to the cell membrane permeability. Notable no significant variation in the antimicrobial activities was found in one way ANOVA at the 95% confidence level (*p*≤0.05), followed by the post hoc Tukey test among the PEF, CHF, and the standard antibiotic, Tetracycline. In the case of gram-negative bacteria, the potency of microbial growth inhibition of PEF and CHF is very much comparable with the tetracycline. These results signify that the potent antimicrobial compounds are mostly extracted through the petroleum ether followed by chloroform and ethyl acetate, which implies their non-polar nature.

Phytochemical analysis indicates that in addition to flavonoids, it may be the alkaloids in phytochemistry which is present in PEF, CHF in a decreasing amount but absent in EAF, though steroid is exclusively present in these three fractions also may responsible too. Phytochemicals exert their antimicrobial effects in different ways; for example, alkaloids intercalate with DNA and inhibit topoisomerase thus blocks DNA synthesis^46^. Another similar work reported where alkaloids of *C. citrinus* showed antibacterial potency against *S. aureus* and *P. areuginosa* through inhibition of ATP-dependant efflux pumps^25^. This pump helps bacteria to prevent the accumulation of drugs or compounds (antibiotics) inside the cells. Another study indicated the essential oils of *C. citrinus* (both leaves and flower) to have *in vitro* antibacterial activity against *B. cereus, E. coli, S. aureus*, and *S. typhi* due to the presence of phenolics and terpenoids (mainly monoterpenoids)^47^. Statistical significant differences are present between vehicle control (DMSO) and all the effective fractions (PEF, CHF and EAF) which nullify the questions of any solvent toxicity.

There are strong positive correlations between consumption of polyphenolics enrich foodstuff and lowering the incidence of degenerative chronic diseases such as cancer, arthritis, renal diseases, heart disease, inflammations, neural disorders like brain dysfunction and eye diseases (cataracts) which are related to oxidative stress^48^. Phenolics and flavonoids are the secondary metabolites, present in almost all plants at various levels and positively correlate each other’s quantity^8^, here too. A positive correlation exists between the reduced incidence of cancer and the regular uptake of phenolics-rich plant products^49^. Correlation analysis indicates linear negative correlations between the total phenolics or flavonoids contents and EC_50_ values of DPPH or FRAP assay and their linear positive correlations (*r* = 0.956 and 0.878, *P* = 0.002 and 0.026 respectively) with the total antioxidant capacity. Thus it was concluded that phenolics and flavonoids may contribute respectively 95.6 and 74.9 % of the total antioxidant capacities. Such a strong positive relationship between this phytochemicals’ quantity and *in vitro* free radical quenching activity of the leaf extract fractions of *Manilkara hexandra* (Roxb.) has already been established^6^. Also, these types of correlations are established between the antioxidant activities and the total phenolics and flavonoids contents of many different plants like *Sonchus arvensis*^50^, *Ampelocissus latifolia*^9^, and *Crinum asiaticum*^8^.

## 5. Conclusion

*Callistemon citrinus* leaf extracts contain various types of phytochemicals which were fractionated through solvent partitioning and polarity gradient. Its successive extract fractions have varying degrees of *in vitro* antioxidant and antibacterial activities. The most polar aqueous fraction of the successive extract fractions exhibited the highest antioxidant activities along with the highest level of total phenolics and flavonoids contents and they were also found to be correlated positively. On the other hand, the most non-polar petroleum ether fractionation showed the highest antibacterial potential and also contains the second-highest flavonoid contents amongst all the extract fractions. Thus, aqueous extract fraction, AQF, from *C. citrinus* leaf may be considered as a good source of natural antioxidants and the petroleum ether extract fraction, PEF, as a natural source of antibacterial agents that might be useful against oxidative stress-related diseases and bacterial infection respectively. Thus, these extract fractions might have therapeutic and industrial importance. A further elaborate investigation is needed to explore the active principle(s), mechanism action, dose optimization, and toxic profile before using them as prophylaxis medication mainly in livestock industries.

## Acknowledgements

The authors acknowledge the financial support of SVMCM The Government of West Bengal as the Swami Vivekananda Scholarship for M.Phil. and Ph.D., CSIR and UGC fellowship and the DST-FIST, DST-PURSE, and UGC-DRS-sponsored infrastructural facilities in the Department of Zoology. Prof. A. Mukherjee has kindly authenticated the plant species. Authors express gratitude to Prof. Soumendranath Chatterjee, Department of Zoology, The University of Burdwan, for extending microbial culture facility and generous supply of bacterial strains.

## Authors Contribution

Poulomi Ghosh: Investigation, data curation, Draft manuscript writing; Souren Goswami: Investigation: Sujit Roy: Investigation; Ria Das: Investigation: Tista Chakraborty: Investigation; Sanjib Ray: Conceptualization, Investigation, data analysis and validation, final manuscript writing and editing.

### List of abbreviations

PEF: Petroleum ether fraction
CHF: Chloroform fraction
EAF: Ethyl acetate fraction
MEF: Methanol fraction
AQF: Aqueous fraction
CAQ: Crude aqueous extract
AAE: Ascorbic acid equivalent
TAE: Tannic acid equivalent
QE: Quercetin equivalent
TFC: Total flavonoids content
TPC: Total phenolics content
DMSO: Dimethyl sulfoxide
SEM: Standard error mean
SEF: Successive extract fractions
DE: Dried extract

